# Increased severity of closed head injury or repetitive subconcussive head impacts enhances post-traumatic headache-like behaviors in a rat model

**DOI:** 10.1101/2020.03.14.979047

**Authors:** Dara Bree, Jennifer Stratton, Dan Levy

**Affiliations:** Dept. of Anesthesia, Critical Care and Pain Medicine, Beth Israel Deaconess Medical Centre, 330 Brookline Ave, Boston, MA, USA 02215; Fax: none.; Harvard Medical School, 330 Brookline Ave, Boston, MA, USA 02215; Teva Biologics, Redwood City, CA, USA

**Keywords:** Posttraumatic headache, concussion, repetitive subconcussive head impacts, cutaneous pain hypersensitivity, Anti-CGRP monoclonal antibody

## Abstract

**Introduction:** Posttraumatic headache (PTH) is one of the most common, debilitating and difficult symptoms to manage after a traumatic head injury. The development of novel therapeutic approaches is nevertheless hampered by the paucity of preclinical models and poor understanding of the mechanisms underlying PTH. To address these shortcomings, we previously characterized the development of PTH-like pain behaviors in rats subjected to a single mild closed head injury using a 250 g weight drop. Here, we conducted a follow-up study to further develop this preclinical model by exploring the development of headache-like pain behaviors in male rats subjected to a single, but more severe head trauma (450 g) as well as following repetitive, subconcussive head impacts (150 g). In addition, we tested whether these behaviors involve peripheral CGRP signaling by testing the effect of systemic anti-CGRP monoclonal antibody (anti-CGRP mAb).

**Methods:** Adult male Sprague Dawley rats (total n=138) were subjected to diffuse closed head injury using a weight-drop device, or a sham procedure. Three injury paradigms were employed: a single hit, using 450 g or 150 g weight drop, and three successive 150 g weight drop events conducted 72 hours apart. Changes in open field activity and development of headache-related cephalic and extracephalic mechanical pain hypersensitivity were assessed up to 42 days post head trauma. Treatment included systemic administration of a mouse anti-calcitonin-gene-related peptide monoclonal antibody (30 mg/kg.).

**Results:** Rats subjected to 450 g closed head injury displayed an acute decrease in rearing and increased thigmotaxis, together with cephalic and extracephalic mechanical pain hypersensitivity that resolved by 6 weeks post-injury. Repetitive subconcussive head impacts using the 150 g weight drop, but not a single event, led to decreased vertical rearing as well as prolonged cephalic and extracephalic mechanical pain hypersensitivity. Early and prolonged anti-CGRP mAb treatment inhibited the development of the cephalic, but not extracephalic pain hypersensitivities in both the severe and repetitive subconcussive head impact models.

**Conclusions:** When compared to the data obtained from male rats in the previous study, a more severe head injury gives rise to a prolonged state of cephalic and extracephalic hyperalgesia. Such enhanced headache-like behaviors also occur following repetitive, subconcussive head impacts. Extended headache-like behaviors following severe and repetitive mild closed head injury are ameliorated by early and prolonged anti-CGRP mAb treatment, suggesting a mechanism linked to peripheral CGRP signaling.

## Introduction

Post-traumatic headache (PTH) is one of the most commonly occurring and disabling sequalae of traumatic head injury, in particular following motor vehicle accidents, falls and sport-related activities (1–3). PTH is defined by the International Classification of Headache Disorders (ICHD-3) as a secondary headache that develops within seven days after the head trauma, or regaining of consciousness following the injury to the head (4). PTH shares similar clinical characteristics with primary headaches, in particular migraine and tension-type headache (TTH) (5–8). However, the extent to which these conditions share common pathophysiological mechanisms that can be targeted using current or novel therapeutic approaches remains unclear, in part due to the small number of studies that employ clinically relevant animal models.

One of the key factors that is thought to influence the propensity to develop PTH is preexisting headache or migraine conditions (3). Yet, many cases of PTH arise de novo following a concussive head trauma (8). We have recently characterized a rat model of de novo PTH that displays numerous headache/migraine-like pain behaviors following a single mild closed head injury (CHI) (9). In particular, we observed the development of cephalic mechanical allodynia that could be ameliorated by systemic treatment with a blocking monoclonal antibody that targets calcitonin-gene-related peptide (Anti-CGRP mAb) (9), suggesting the involvement of peripheral CGRP signaling.

Although our rat model as well as other recently developed mouse models (10, 11) are likely to be provide important insights into the mechanisms underlying PTH, they do not fully capture the factors and conditions that lead to PTH as they are limited to the effects of a single mild concussive head trauma. An important factor that may influence the propensity to develop of PTH is the severity of the head trauma. PTH has been suggested to be more common after a mild trauma when compared to more severe events (3). However, moderate and severe traumas are also associated with high prevalence of PTH (12). Nonetheless, some studies suggest either no correlation between the severity of the trauma and incidence of PTH (13, 14), increased frequency of persistent and more severe PTH following moderate or severe head trauma (15), and similar sensory abnormalities regardless of the injury severity (7). Although acute head trauma is considered the leading cause of PTH, repetitive subconcussive head impacts, such as those occurring during professional and recreational sporting events, and which are difficult to detect or quantify, can also lead to post-traumatic neurological sequelae, and potentially PTH (16). Here, we aimed to address these additional factors, and further develop a clinically relevant PTH rat model. We first characterized time-course changes in cephalic and extracephalic pain sensitivity in a model that mimics a more severe head trauma. We then examined the cumulative effect of repetitive subconcussive head impacts on the development of these pain behaviors. Finally, we addressed the relative contribution of peripheral CGRP signaling in these two new PTH modeling paradigms by studying the analgesic effect of anti-CGRP mAb.

## Materials & Methods

### Animals

All experiments were approved and conducted in compliance with the institutional Animal Care and Use Committee of the Beth Israel Deaconess Medical Centre, and the ARRIVE (Animal Research: Reporting of *in vivo* Experiments) guideline. Subjects were adult male Sprague-Dawley rats (total n=138). Animals were obtained from Taconic, USA, and were 8-9 weeks at arrival. They were housed in pairs with food and water available *ad libitum* under a constant 12-hour light/dark cycle (lights on at 7:00 am) at room temperature. Studies were initialized after a week of acclimatization in the vivarium. All procedures and testing were conducted during the light phase of the cycle (8:00 am to 3:00 pm). Experimental animals were randomly assigned to either sham or CHI as well as to the different pharmacological treatment groups.

### Experimental Closed Head Injury (CHI) models

We have previously used a mild CHI model involving a 250 g weight drop. Here, we tested 3 different CHI paradigms, employed in separate cohorts of rats. The first paradigm involved a single, more severe hit, using a 450 g weight drop. The second paradigm employed a single 150 g weight drop. The third paradigm involved 3 successive 150 g weight drop events, conducted 72 hours apart. All head injury paradigms were conducted while the animals were anesthetized with 3% isoflurane and placed chest down directly under the weight-drop concussive head trauma device. The device consisted of a hollow cylindrical tube (80 cm) placed vertically over the center of the rat’s head. To ensure consistency of the hit location, animals were placed under the weight drop apparatus so that the weight struck the scalp slightly anterior to the center point between the ears. A foam sponge (thickness 3.81 cm, density 1.1 g/cm^3^) was placed under the animals to support the head while allowing some linear anterior-posterior motion without any angular rotational movement at the moment of impact. A repeated strike was prevented by capturing the weight after the first strike. Surviving animals regained their righting reflex within 2 minutes (which likely reflects the recovery from anesthesia), and were returned immediately to their home cages for recovery. Animals were assessed during the first 7 days post injury for any behavioral abnormalities suggestive of a major neurological (i.e. motor) impairment. For all paradigms, sham animals were anesthetized for the same period of time as CHI animals, but not subjected to the weight drop.

### Open field behavior

Changes in locomotor activity, exploratory behavior, and anxiety-like behavior in an open field arena (43 × 43 × 30 cm) were monitored as previously described using Activity Monitor (Med Associates, Vermont, USA). The monitoring system records the movement of animals in the horizontal (X-Y axis) and vertical (Z-axis) planes using 16 infrared beams and detectors, spaced 2.54 cm apart. Data analyzed included total distance moved, vertical rearing events, and relative time spent in the center of the arena (30 cm X 30 cm) during a 20-minutes session. The arena was lit with a single white LED bulb on a dimmer switch to maintain a homogenous lighting across the arenas (80 lux). The arena was cleaned with a mild detergent and dried to remove odor cues between successive rats. Given that activity in open field testing is based on novelty exploration, and repeated exposure decreases novelty, testing was limited to 4 trials: one baseline trial, and 3 post injury trials, with the last one conducted on day 14 post injury.

### Assessment of tactile pain hypersensitivity

To assess changes in tactile pain sensitivity, we employed a method that was previously used by us and others to study PTH- and migraine-related pain behaviors (9, 17–19). Briefly, animals were placed in a transparent acrylic tube (20.4 cm × 8.5 cm) closed on both ends. The apparatus was large enough to enable to the animals to escape the stimulus. Animals were habituated to the arena for 15 minutes prior to the initial testing. In order to determine if the animals developed pericranial (cephalic) tactile hypersensitivity, the skin region, including the midline area above the eyes and 2 cm posterior, was stimulated with different von Frey (VF) filaments, using an ascending stimulus paradigm (0.6 g–15 g/force, 18011 Semmes-Weinstein Anesthesiometer kit). Development of hind paw tactile hypersensitivity was tested by stimulating the mid-dorsal part of the hind paw using the VF filaments. We evaluated changes in withdrawal thresholds, as well as non-reflexive pain responses to the stimulation using a method previously described in this and other headache models (20–22) by recording 4 behavioral responses adapted from Vos et al. (23) as follows: 0) *No response*: rat did not display any response to stimulation 1) *Detection*: rat turned its head towards stimulating object and latter is explored usually by sniffing; 2) *Withdrawal*: rat turned its head away or pulled it briskly away from stimulating object (usually followed by scratching or grooming of stimulated region); 3) *Escape/Attack*: rat turned its body briskly in the holding apparatus in order to escape stimulation or attacked (biting and grabbing movements) the stimulating object. Starting with the lowest force, each filament was applied 3 times with an intra-application interval of 5 seconds and the behavior that was observed at least twice was recorded. For statistical analysis, the score recorded was based on the most aversive behavior noted. The force that elicited two consecutive withdrawal responses was considered as threshold. To evaluate pain behavior in addition to changes in threshold, for each rat, at each time point, a cumulative pain score was determined by combining the individual scores (0–3) for each one of the VF filaments tested as in Vos et al (23). All tests were conducted and evaluated in a blinded manner.

### Pharmacological treatments

Anti-CGRP mAb and its corresponding isotype IgG were provided by TEVA Pharmaceuticals, and formulated in phosphate buffered saline (PBS). mAb and IgG were injected i.p at a dose of 30 mg/kg, immediately after the head injury and every 6 days subsequently. This dosing regimen has been shown previously to alleviate PTH-like pain behaviors (i.e. cephalic allodynia) following mild CHI in male rats (9, 22), and male mice (11), as well as pain behaviors in other chronic migraine models (24).

### Data analyses

Group size (n=8-15) were determined using *a priori* power analysis (G*Power 3.1 software). Calculations were based on our previous and pilot data showing that they were sufficient to detect large (*d* > 0.8) effect sizes in at least 80% power, and α = 0.05. Statistical analyses of experimental data were conducted using Graph pad prism (version 8.0). All data are presented as the means±standard error of the mean. Mechanical pain threshold data, obtained using VF filaments, was log-transformed, which is more consistent with a normal distribution (25). Changes in open-field behavior, and behavioral responses to mechanical stimuli were analyzed using data from sham and CHI animals using a mixed design ANOVA to determine the effects of time and treatment. The same analysis was used to determine the effect of anti-CGRP mAb vs the corresponding IgG control. All data included passed the Brown-Forsythe test, indicating equal variance. We used Fisher’s LSD post hoc tests, and correction for multiple comparisons (Type I error) was conducted using the Benjamini and Hochberg false discovery rate (FDR) controlling procedures

## Results

### The effects of 450 g weight drop injury

#### Open field behavior

In our previous studies, we examined changes in pain behaviors resembling PTH in response to a mild, concussive CHI involving a single 250 g weight drop in rats (9, 22). In this CHI paradigm, male rats displayed acute changes in open field behavior, as well as mechanical pain hypersensitivity, limited to the cephalic region, that recovered at day 14 post CHI. Here we asked whether a more severe CHI paradigm, involving a 450 g weight drop, gives rise to more robust changes in the open field testing as well as a more pronounced pain phenotype. In our earlier studies, all animals subjected to the 250 g weight drop protocol recovered. Here, however, we observed a higher mortality rate in animals subjected to a single 450 g weight drop (~20% of the animals did not survive the CHI), suggesting a more severe head (and likely brain) trauma. In open field testing, when compared to sham, animals subjected to the 450 g weight drop injury did not exhibit changes in total distance travel (Time F_(3, 42)_ = 2.9, p<0.07; Treatment F_(1, 14)_ = 1.1, p = 0.3, Figure 1(**b**)). However, a prolonged decrease in exploratory rearing activity was not noted at 3-14 days post the CHI (Time F_(3, 42)_ =2.8, p<0.05: Treatment F_(1, 14)_ = 24.5, p<0.001; q<0.001, Figure 1(**c**)). In addition, head injured animals displayed an acute decrease in center zone exploration at 3 days post CHI ((Time F_(3, 42)_ = 4.5, p<0.001: Treatment F_(1, 14)_ = 4.7, p < 0.05; q<0.001; Figure 1(**d**)), suggesting elevated anxiety levels.

**Figure 1:**
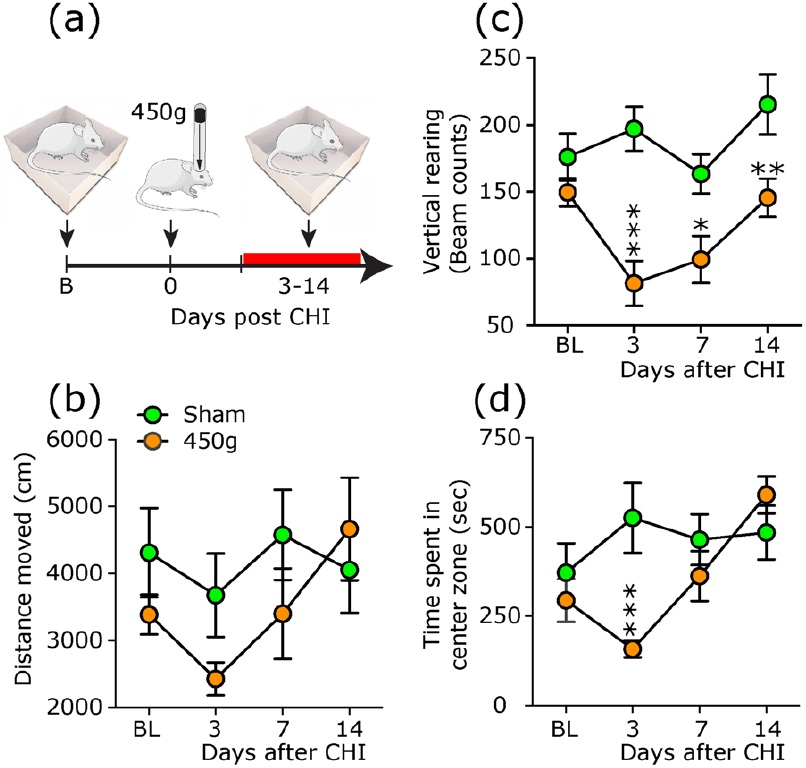
Changes in open-field behaviors in animals subjected to a CHI involving a 450 g weight drop or a sham procedure. (a) Scheme of the experimental design. Rats were subjected to a baseline open-field testing, following by CHI, and then additional testing 3-14 days later. Changes in distance moved (b), exploratory vertical rearing (c), and time spent in the center zone (d). Mixed design ANOVA, followed by post-hoc test between CHI and sham animals indicate a decrease in vertical rearing at 3-14 post CHI, and acute decrease in time spent in center zone (indicating increased thigmotaxis) at 3 days following CHI. Data are means±SEM (sham: n=8; CHI: n=8). * q< 0.05; ** q<0.01, *** q<0.001 (FDR-corrected values after Fisher’s least significant difference post-hoc test at selected time points vs similar time points in sham control animals). FDR, false discovery rate; CHI, mild closed-head injury.

#### Tactile pain sensitivity

Sensory testing revealed prolonged cephalic hyperalgesia when compared to sham animals, including decreased pain threshold, and increased pain scores that were maximal at 3 days post CHI and recovered only by day 42 (Threshold: Time F_(5, 102)_ = 7.2, p<0.001; Treatment F_(1, 24)_= 50.3, p<0.0001; q<0.001 for 3-30 day time points all time points; Pain score; Time F_(5, 102)_ = 4.2, p<0.05; Treatment F_(1, 26)_= 13.0, p <0.01 for days 3 and 30; q<0.05 for days 7 and 14; Figures 2(**b**) and (**c**)). In addition to the pronounced cephalic mechanical pain hypersensitivity, head injured rats also displayed a delayed extracephalic mechanical pain hypersensitivity that was manifested as decreased hind paw pain threshold on days 14-42 post CHI (Time; F_(5, 102)_ = 3.5, p<0.01: Treatment F_(1, 24)_ = 15.2, p <0.001; q<0.001 for days 14, 30 and 42), and increased pain score on days 30 and 42 post CHI (Time F_(5, 102)_ = 5.9, p<0.01: Treatment F_(1, 24)_ = 4.1, p <0.05; q<0.001 for days 30 and 42; Figure 2(**d**) and (**e**)).

**Figure 2:**
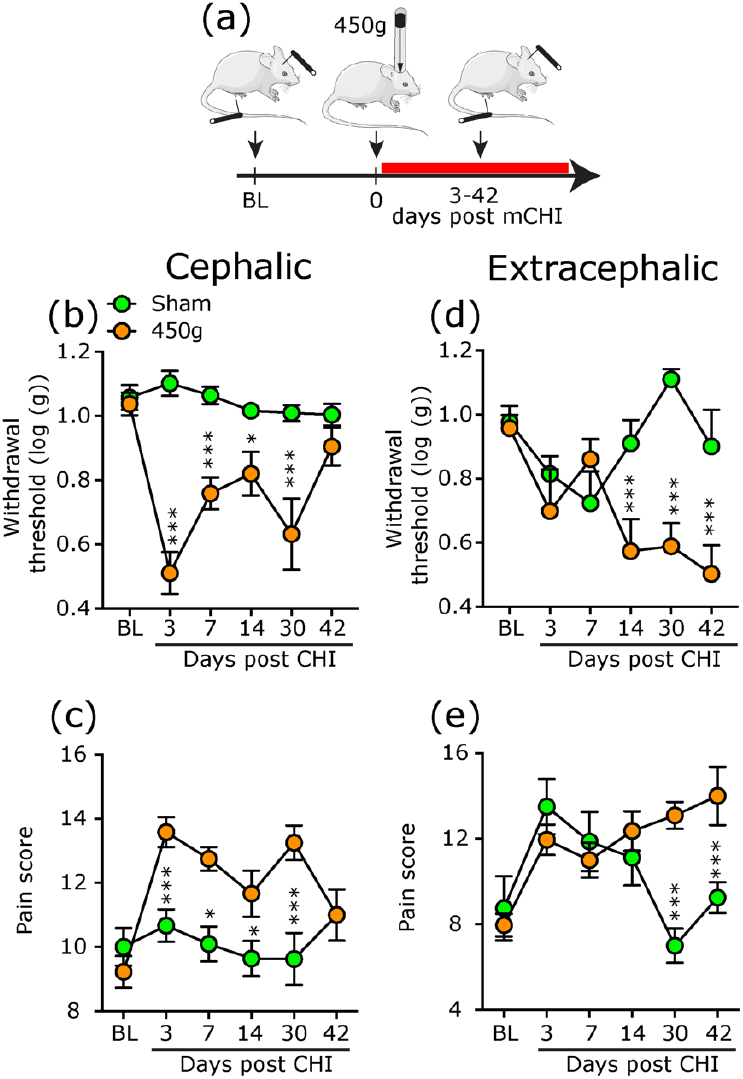
Development of prolonged cephalic and extracephalic cutaneous mechanical hypersensitivity following a 450 g weight drop CHI. (a) Schematic of the experimental design. Rats underwent baseline von Frey testing of mechanical pain sensitivity at the cephalic and extracephalic (hind paw) regions, followed by CHI, and further nociceptive testing at these locations 3-42 days later. Time course changes in cephalic (b) and extracephalic (d) mechanical pain withdrawal thresholds and corresponding cumulative pain scores at the cephalic (c) and extracephalic (e) regions. Mixed-design ANOVA, followed by post-hoc test between CHI and sham animals indicate a prolonged decrease in withdrawal thresholds and increase in pain scores following CHI at the cephalic, and extracephalic regions. Data are means±SEM (Sham: n=15; mCHI: n=11). * q<0.05, *** q<0.001 (FDR-corrected values after Fisher’s least significant difference post-hoc test at selected time points vs similar time points in sham control animals).

### The effects of repetitive subconcussive 150 g weight drop head impacts

#### Open field testing

To address the behavioral consequences of subconcussive head impacts, we initially tested the effects of a milder head trauma, involving a single 150 g weight drop. In open field testing, when compared to sham treated animals, animals subjected to a 150 g mild CHI did not display significant time-dependent changes in any of the 3 parameters tested (distance moved: Time F_(3, 42)_ = 5.0, p<0.05; Treatment F_(1, 14)_ = 4.3; P=0.06; Rearing; Time F_(3, 42)_ = 0.8, p = .5; Treatment F_(1, 14)_ = 3.1, p<0.1; Center zone time: Time F_(3, 42)_ = 0.6, p<0.6: Treatment F_(1, 14)_ = 2.2; P=0.16; Figure 3 (**b**), (**c**), and (**d**)). We next asked whether repetitive, mild CHI have a cumulative effect on the open field behavior. Three consecutive 150 g weight drops, conducted 72 hours apart, did not affect the distance moved (Time F_(3, 42)_ = 3.0, p<0.05; Treatment F_(1, 14)_ = 1.7; p=0.2; Figure 3(**f**)), but led to a time-dependent decrease in rearing (Time F_(3, 42)_ = 5.5, p<0.001; Treatment F_(1, 14)_ = 20.5, p<001; q<0.001 for days 2,7 14 post injury, sham vs repetitive mild CHI; Figure 3(**g**)), indicating a concussion related response (26). Animals subjected to repetitive mild CHI did not display any changes in center zone activity (Time F_(3, 42)_ = 1.3, p=0.3; Treatment F_(1, 14)_ = 3.9; p=0.07; Figure 3(**h**)), indicating no increase in anxiety level.

**Figure 3:**
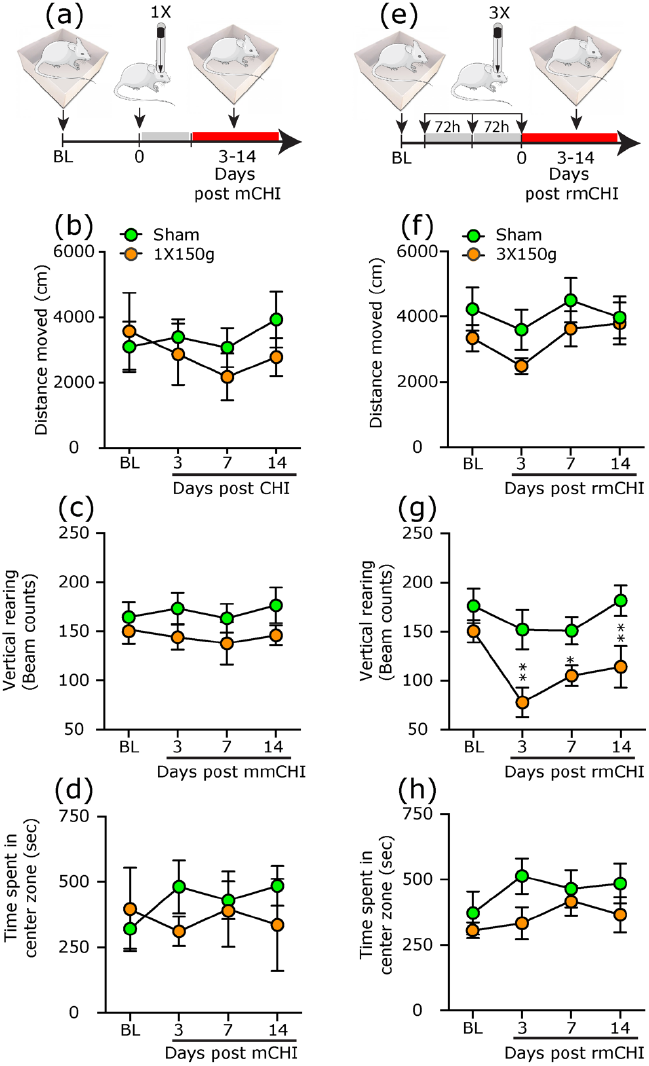
Changes in open-field behaviors in animals subjected to a single mild closed head impact (mCHI), or repetitive impacts (rmCHI) using 150 g weight drop. (a) Scheme of the experimental design for the 1X150 g mCHI model. Rats were subjected to a baseline open-field testing, following by mCHI, and additional testing 3-14 days later. (b) total distance moved, (c) exploratory vertical rearing, and (d) time spent in center zone (means±SEM; sham: n=8; rCHI: n=8). (e-h) Scheme of the experimental design for the 3X150 g rmCHI model and related open-field testing (means±SEM; sham: n=8; rmCHI: n=8). Mixed design ANOVA, followed by post-hoc test between CHI and sham animals indicate an acute decrease in vertical rearing at 3-14 days following a rmCHI. * q< 0.05; ** q<0.01 (FDR-corrected values after Fisher’s least significant difference post-hoc test at selected time points vs similar time points in sham control animals). FDR, false discovery rate; CHI, mild closed-head injury.

#### Tactile pain sensitivity

Given the open field data suggesting that a single mild CHI involving a 150 g weight drop is subconcussive, we next examined whether this head trauma paradigm leads to increased nociceptive behavior. We then assessed changes in pain responses following the repetitive mild CHI paradigm. When compared to sham treatment, animals subjected to a single 150 g mild CHI did not show any changes in cephalic mechanical pain threshold (Time F_(3,54)_ = 3.3, p<0.05: Treatment F_(1, 22)_ = 0.04; P=0.83; Figure 4(**b**)), or pain score (Time F_(3, 54)_ = 3.1, p<0.05: Treatment F_(1, 22)_ = 0.09; P=0.8; Figure 4(**c**)). Repetitive mild CHI however led to a prolonged, and time-dependent mechanical pain hypersensitivity. This hyperalgesic response was manifested as a decrease in threshold that was already maximal at 3 days following the last head injury and resolved by day 42 (Time F_(5, 110)_ = 6.7, p<0.001; Treatment F_(1, 26)_ = 16.4, p<0.001; q<0.01 for days 3-30, sham vs repetitive mild CHI, Figure 4(**e**)) and a similar profile of increased pain score (Time F_(5, 104)_ = 3.3, p<0.01; Treatment F_(1, 26)_ = 11.0, p<0.01, q<0.001 for days 3-14, q<0.01 for day 30, sham vs repetitive mild CHI; Figure 4(**f**)).

**Figure 4:**
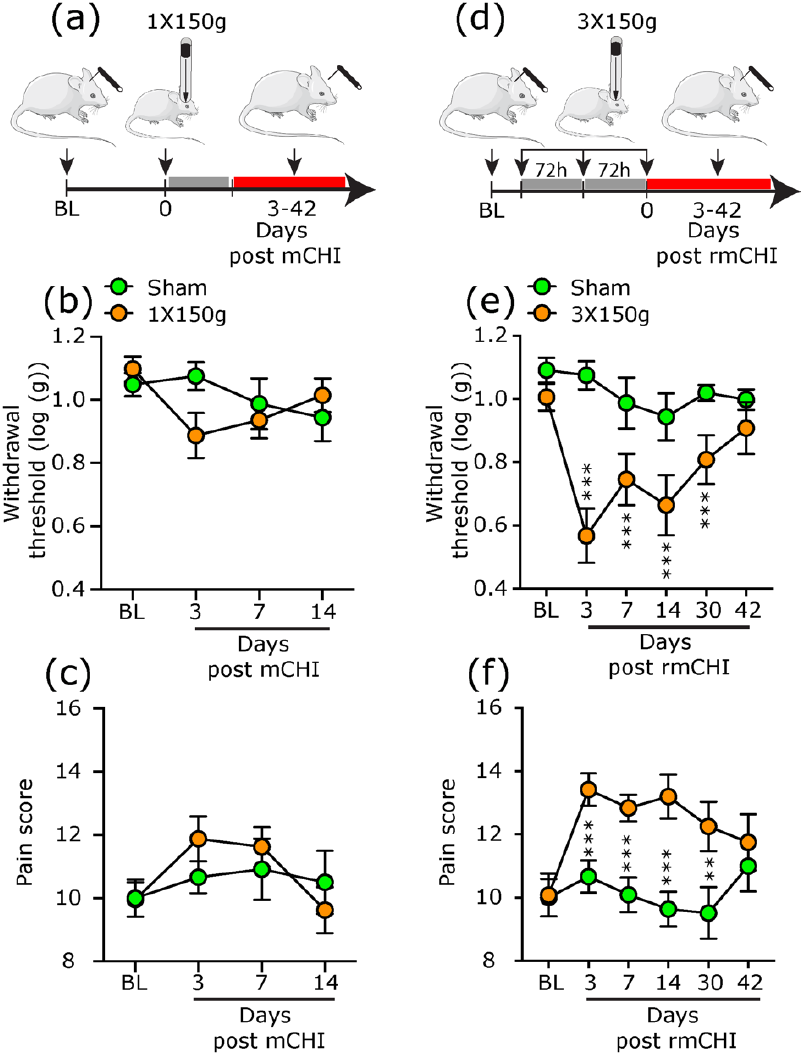
Development of prolonged cephalic cutaneous mechanical hypersensitivity following rmCHI. Schematic of the experimental design. Rats underwent baseline von Frey testing of mechanical pain sensitivity at the cephalic region, followed by either a single 150 g mCHI (a), or 3X150 rmCHI (d) and further nociceptive testing 3-42 days later. Time course changes in cephalic mechanical pain withdrawal thresholds (b) and corresponding cumulative pain scores (c) following a single mCHI (means±SEM; Sham: n=8; CHI: n=8). Changes in threshold (e) and pain score (f) in response to rmCHI (means±SEM; Sham: n=8; rmCHI: n=8). Mixed design ANOVA, followed by post-hoc tests between CHI and sham animals indicate a prolonged decrease in withdrawal thresholds and increase in pain score 3-30 days following rmCHI. ** q<0.01, *** q<0.001 (FDR-corrected values after Fisher’s least significant difference post-hoc test at selected time points vs similar time points in sham control animals).

Assessment of changes in extracephalic pain behavior revealed similar effects. A single 150 g mild CHI did not affect extracephalic mechanical pain sensitivity (Threshold: Time F_(3, 46)_ = 4.0, p<0.05; Treatment F_(1, 18)_ = 0.1; P=0.8; Pain score: Time F_(3, 46)_ = 1.5, p < 0.05; Treatment F_(1, 22)_ = 0.3, p=0.5; Figure 5(**b**) and (**c**)). However, animals subjected to repetitive subconcussive mild closed head impacts exhibited extracephalic mechanical pain hypersensitivity manifested as a decrease in pain thresholds that developed with a delay and was significant on days 14 and 30 (Time F_(5, 110)_ = 3.6, p<0.01; Treatment F_(1, 26)_ = 5.5, p<0.05; q<0.001 for both, sham vs repetitive mild CHI; Figure 5(**e**)).

**Figure 5:**
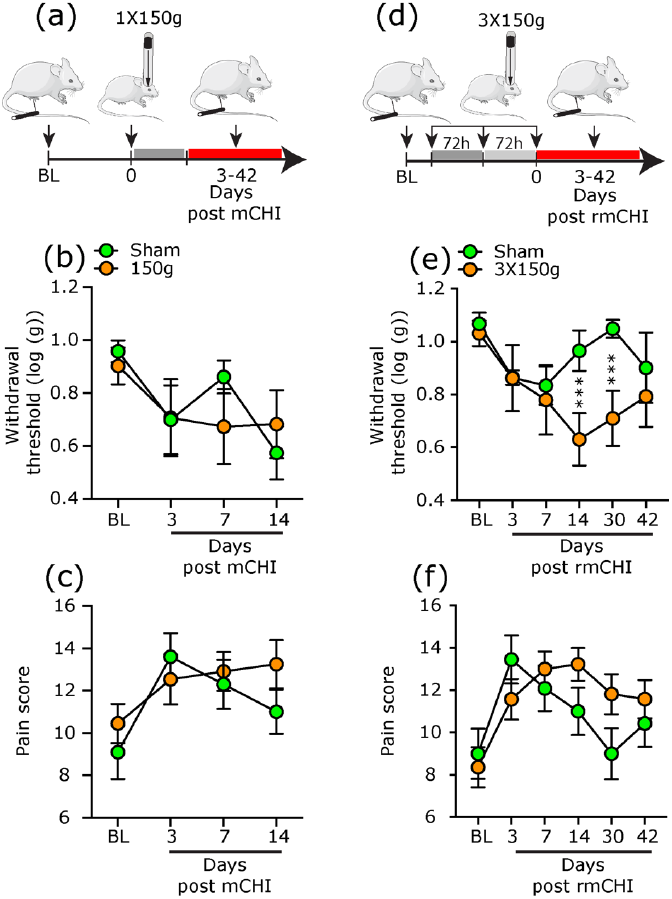
Development of delayed extracephalic cutaneous mechanical hypersensitivity following rmCHI. Schematic of the experimental design. Rats underwent baseline von Frey testing of mechanical pain sensitivity at the hind paw, followed by either a single 150 g mCHI (a), or 3X150 g rmCHI (d) and further nociceptive testing 3-42 days later. Time course changes in hind paw mechanical pain withdrawal thresholds (b) and corresponding cumulative pain scores (c) following a single mCHI (means±SEM; Sham: n=8; CHI: n=8). Changes in threshold (e) and pain score (f) in response to rmCHI (means±SEM; Sham: n=8; CHI: n=8). Mixed design ANOVA, followed by post-hoc test between CHI and sham animals indicate a delayed decrease in withdrawal thresholds 14-30 days following rmCHI. Data are means±SEM. *** q<0.001 (FDR-corrected values after Fisher’s least significant difference post-hoc test at selected time points vs similar time points in sham control animals).

### The effect of systemic anti-CGRP mAb treatment

In our previous study of male rats subjected to a 250 g weight drop CHI, systemic administration of a mAb that blocks CGRP function ameliorated the cephalic mechanical pain hypersensivity (9). Here, a similar mAb treatment was also effective in ameliorating the prolonged cephalic pain hypersensitivity that developed in rats subjected to acute 450 g weight drop CHI. As Figures 6 (**b**) and (**c**) depict, there was a statistically significant difference between the effects of the control IgG and the anti-CGRP mAb with regard to threshold changes (Time F_(5,70)_ = 7.4, p<0.0001; Treatment F_(1,14)_ = 9.6, p<0.01). Post hoc analysis revealed a prolonged decrease in cephalic pain thresholds in the IgG treatment group (q<0.01 for days 3-14, q<0.05 for day 30), but not in the anti-CGRP mAb treatment group (q>0.05 for all time points). Similarly, the two treatment groups were statistically different with regard to pain scores (Time F_(5,70)_ = 4.4, p<0.01; Treatment F_(1,14)_ = 4.9, p<0.05). Post hoc analysis revealed a pronged increased in cephalic pain score in IgG-treated animals (q<0.01 for days 3-30), but no change animals treated with the anti-CGRP mAb (q>0.05 for all time points). There was no difference between the IgG and anti-CGRP mAb treatments with regard to the extracephalic pain hypersensitivity, including for threshold changes (Time F_(5,70)_ = 2.5, p<0.05; Treatment F_(1,14)_ = 0.11, p=0.9; Figure 6(**d**)) and for pain score (Time F_(5,70)_ = 0.4, p = 0.9; Treatment F_(1,14)_ = 0.1, p = 0.9; Figure 6(**e**)).

**Figure 6:**
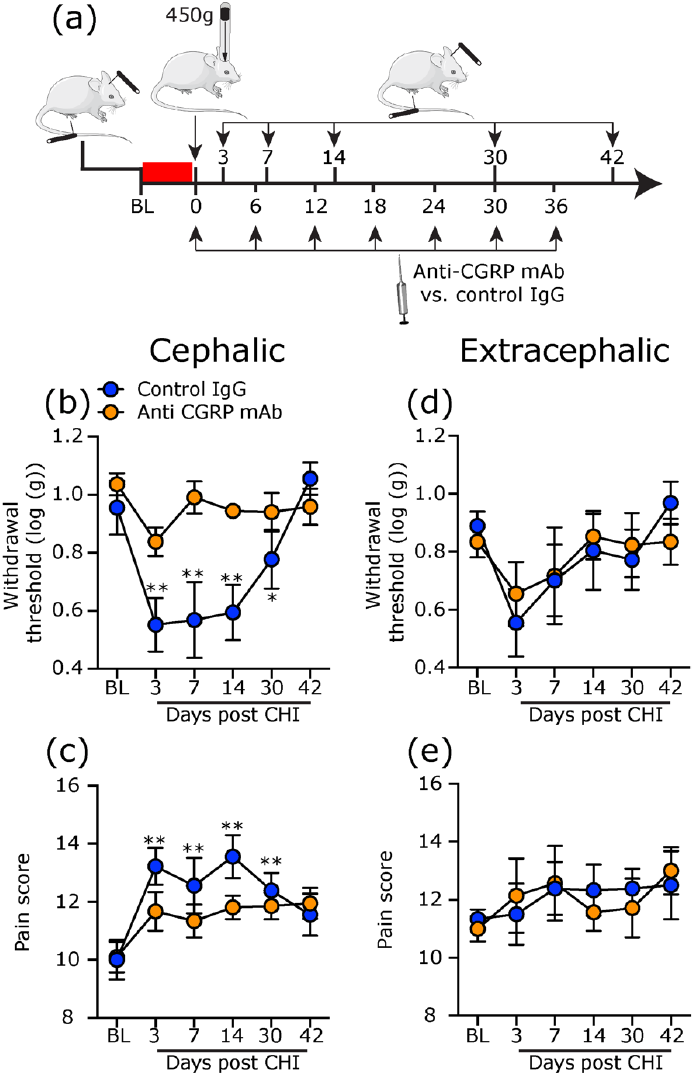
Early and prolonged treatment with anti-CGRP mAb ameliorates cephalic, but not extracephalic pain hypersensivity in rats subjected to 450 g CHI. Schematic of experimental design. Rats underwent baseline von Frey testing of cephalic and extracephalic mechanical pain sensitivity, followed by CHI. Anti-CGRP mAb or a control IgG were administered immediately after the CHI and then every 6 days. Mixed design ANOVA revealed effects of treatment on the decrease in cephalic withdrawal threshold (b), and the increase in pain score (c), but not on the extracephalic threshold (d) or pain score (e). Means±SEM (Anti-CGRP: n=8; control IgG: n=8). * q<0.05; ** q<0.01 (FDR-corrected values after Fisher’s least significant difference post-hoc test for post CHI values vs baseline).

Treatment with anti-CGRP mAb also exerted an anti-nociceptive in rats subjected to repetitive subconcussive 150 g weight drop head impacts. As Figures 7(**b**) and (**c**) depict, there was a statistically significant difference between the IgG and anti-CGRP mAb treatment groups with regard to threshold changes (Time F_(5,70)_ = 8.5, p<0.001; Treatment F_(1,14)_ = 4.7, p<0.05) and pain scores (Time F_(5,70)_ = 4.0, p<0.01; Treatment F_(1,14)_ = 5.1, p<0.05). Post hoc analysis revealed a decrease in cephalic pain thresholds in IgG-treated animals (q<0.01 for days 7-30), but no change in animals treated with the anti-CGRP mAb (q>0.05 for all time points). Similarly, there was an increase in cephalic pain score in the IgG treatment group (q<0.01 for days 7-14; q<0.05 for day 30), but no change in the anti-CGRP mAb treatment group (q>0.05 for all time points). There was no difference between the IgG and anti-CGRP mAb treatments with regard to the extracephalic pain hypersensitivity including changes in threshold (Time F_(5,70)_ = 3.0, p<0.05; Treatment F_(1,14)_ = 1.4, p=0.24; Figure 7(**d**), as well as in pain score (Time F_(5,70)_ = 2.4, p<0.05; Treatment F_(1,14)_ = 0.7, p = 0.39; Figure 7(**e**)).

**Figure 7:**
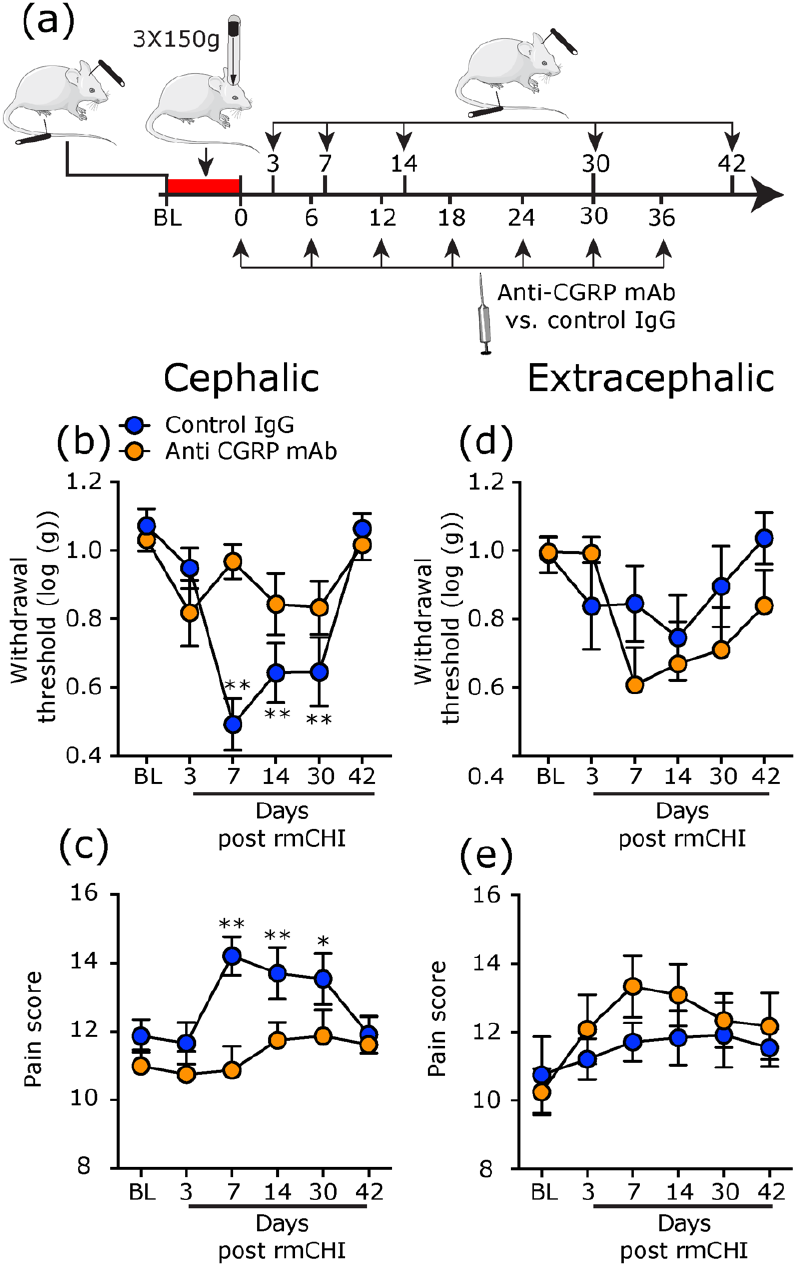
Early and prolonged treatment with anti-CGRP mAb ameliorates cephalic, but not extracephalic pain hypersensivity in rats subjected to rmCHI. Schematic of experimental design. Rats underwent baseline von Frey testing of cephalic and extracephalic mechanical pain sensitivity, followed by rmCHI. Anti-CGRP mAb or a control IgG were administered immediately after the last weight drop, and then every 6 days. Mixed design ANOVA revealed effects of treatment on the decrease in cephalic withdrawal threshold (b), and the increase in pain score (c), but not on the extracephalic threshold (d) or pain score (e). Means±SEM (Anti-CGRP: n=8; control IgG: n=8). ** q<0.01 (FDR-corrected values after Fisher’s least significant difference post-hoc test for post mild CHI values vs baseline).

## Discussion

In the present study, we have extended the characterization of a preclinical rat model of PTH, using additional closed head trauma paradigms to investigate the consequences of a more severe head injury, and repetitive subconcussive events. Our data suggests that both paradigms are associated with increased behavioral symptoms suggestive of traumatic brain injury and prolonged pain hypersensivity when compared to our original mild head injury paradigm (9).

In our previous rat studies, we have used a model of diffuse closed head injury by employing a 250 g weight drop from 80 cm height. Here, to produce a more severe CHI we increased the gravity force by employing a heavier mass (450 g weight), and maintained the same height as in our previous CHI paradigm. The finding that the 450 g CHI paradigm was associated with a ~20% mortality rate, while our earlier studies indicated no mortality using the milder 250 g CHI paradigm (9, 22) points to increased injury severity, and is in agreement with previous studies that investigated the effects of a graded mechanical impact levels (27). Our open-field testing data indicated no change in total distance traveled suggesting no impairment in gross motor activity, similar to the effect of a 250 g weight drop injury. However, animals subjected to the 450 g weight drop injury exhibited decrease in exploratory rearing activity that lasted much longer than that observed in animals subjected to the milder (250 g) CHI paradigm. In the context of this injury model, decreased rearing is likely to indicate traumatic brain injury (26) and possibly also a related increase in anxiety level (28). Furthermore, rats subjected to the 450 g CHI also displayed increased thigmotaxis, another measurement of increased anxiety levels in open field testing (28), a behavior which was not observed previously in the milder CHI paradigm. The prolonged reduction in rearing activity following the more severe head injury could also reflect migraine-like head pain as it was also observed following noxious stimulation of the cranial meninges (19), as well as in response to peripheral and systemic administration of the headache trigger CGRP (29).

In our previous studies, male rats subjected to 250 g weight drop head injury protocol displayed a gradual increase in cephalic mechanical pain sensitivity that reached peak values at 7 days and resolved a week later. Here, increasing the severity of the head injury, using the 450 g weight drop, gave rise to an earlier onset of cephalic pain hypersensitivity, that was already maximal on day 3 post injury, and lasted for more than 4 weeks. Rats subjected to this more severe CHI protocol also displayed delayed extracephalic pain hypersensitivity. The rapid development of the cephalic pain hypersensitivity in this CHI paradigm could be mediated by an acute trauma endured by deep cranial tissues, in particular the calvarial periosteum and intracranial meninges, and the ensuing sensitization of their trigeminal sensory innervation (21, 30). It is also possible that the acute increase in anxiety levels noted on day 3 post injury also contributed to the rapid increase in nociceptive behavior. The mechanisms underlying the prolongation of the enhanced trigeminal nociceptive response may involve sustained peripheral tissue damage that maintains the sensitized state of the deep cranial nociceptors (e.g. sterile inflammation), or the second order nociceptive neuron in the medullary dorsal horn that receive their input. Alternatively, damage to modulatory pain pathway, including ascending and/or descending pathways could also play a role (31). The delayed development of the extracephalic pain hypersensitivity and its persistent nature supports this notion and further suggest the involvement of sensory thalamic nuclei, similar to the extended allodynia observed following stimulation of meningeal afferents and during a migraine attack (32, 33).

Prospective studies suggest that the prevalence of persistent PTH is inversely correlated with the severity of the head trauma and the resultant brain injury, as is the case with the frequency of other TBI-related pain syndromes (34). It is not clear however whether these differences are due to biological factors, or the result of reporting bias. While persistent PTH is clearly a common sequalae of moderate and severe TBI (15), it is also unclear whether its clinical manifestations are less severe than in PTH cases that develop following milder traumas. Our study, which assessed pain hypersensitivity as a surrogate measure of headache, points to an enhanced pain phenotype following a more severe head trauma, when compared to our previous data using the milder weight drop head injury model. Further studies will be required to determine whether the 450 g weight drop injury paradigm we employed herein inflicts peripheral and/or brain damage that are within the parameters of those considered to occur following a milder concussive head/brain trauma or more severe ones. Nonetheless, if the 450 g weight drop injury model can be equated to a moderate/severe TBI, our data may explain the finding that subjects with PTH following these types of TBI are more likely to experience higher frequency of headaches (several per week/daily) than those with PTH following a milder trauma (1, 15).

Repetitive concussive head injuries, as well as subconcussive events, can lead to numerous chronic neurobehavioral problems including persistent headache (35). Although there is increased interest in the pathophysiological outcomes of repetitive mild TBI, including their role in PTH (36), the contribution of subconcussive head impacts to headache remains poorly studied. Here, we first established a subconcussive weight drop paradigm, by showing that a single 150 g weight drop does not exhibit any changes in open field activity. We further observed that this acute subconcussive impact does not lead to headache-like pain behaviors, suggesting a lack of acute cephalic injury. A key finding, however, was that repeating these subconcussive events gave rise to persistent deficit in open field activity, which likely implicates mild to moderate brain injury. In addition, we observed persistent increases in cephalic and extracephalic pain sensitivity, resembling those observed in animals subjected to the 450 g. weight drop injury. We modeled the repetitive head injury paradigm to include 72 hours of recovery time, which mimics regular events in professional as well as in many amateur contact sports, such as American football, boxing, and hockey. It will be of interest to examine in future studies the minimal duration of recovery time, as well as number of repetitive events, that may be considered safe and do not affect open field and pain behaviors. Examining these factors can help future studies that address issues such as return to play following sport related head injuries. The mechanisms by which repetitive, subconcussive head impacts gives rise to persistent headache-like pain behavior could potentially implicate cumulative effects of the acute subconcussive impacts. The similarity to the persistent headache-like behavioral changes observed in an animal model of repeated administration of inflammatory mediators to the cranial meninges (17), or systemic nitroglycerin (37) points to the possibility that repeated subconcussive head impacts engage pathophysiological mechanisms similar to those that perpetuate persistent migraine pain. Whether the enhanced pain behaviors are due to cumulative effects of meningeal irritation and related inflammation, or brain injury will need to be addressed in future studies. We would also like to entertain the possibility that local or remote changes evoked by the initial head impacts promote peripheral and/or central hyperalgesic priming (38–40) that gives rise to a prolonged nociceptive response following an additional trauma event.

Systemic repeated administration of the anti-CGRP mAb, starting immediately after the acute 450 weight drop injury, or the last subconcussive 150 g head impact was able to inhibit the increased cephalic pain hypersensivity, when compared to treatment with the control IgG. The anti-nociceptive response of the anti-CGRP mAb was overall similar to that previously observed in the 250 g weight drop injury model in male rats with one exception: that the treatment was already effective on day 3 in the 450 g impact model, but started to become effective only at day 7 in the repetitive subconcussive model. The effectiveness of the mAb therapy in these PTH models implicates a peripheral mechanism linked to CGRP signalling, potentially related to meningeal and/or periosteal neurogenic inflammation. The finding that anti-CGRP treatment did not mitigate the extracephalic pain hypersensitivity in both head trauma models, points to the possibility that additional mechanisms related to CNS plasticity or injury are also involved in the mechanism underlying the extracephalic pain hypersensitivity in PTH.

An important limitation of the current study is the lack of female subjects. Females may be more susceptible to developing persistent PTH (8, 41), and our recent study points to an enhanced PTH-like behaviors in female rats in the 250 g CHI paradigm (42). Whether females develop PTH-like symptoms in the new CHI paradigms we tested herein beyond those observed in males will have to be examined in future studies. Because anti-CGRP mAb treatments is less effective in female rats in the 250 g CHI model (when compared to males) (42), it will also be of interest to examine in future studies whether these CGRP-related sex differences also present in the more severe closed-head injury or repetitive subconcussive closed head impact models.

## Conclusion

We characterized two additional preclinical traumatic closed head injury paradigms that can be used to study the pathophysiology of PTH and its treatment. Our injury paradigms employing a single 450 g weight drop, or repetitive 150 g subconcussive head impacts provide a platform to study more persistent PTH-like behaviors. The repetitive head injury paradigm may also be used to investigate the origin of PTH in sport related injuries. Our data further support the notion that the mechanisms underlying the persistent cephalic pain hypersensitivity in PTH in males involve peripheral CGRP signaling, while those that mediate extracephalic pain hypersensitivity are CGRP-independent.

## Acknowledgments

Dr. Levy is supported by NIH Grants R01NS078263; R01NS086830, R21NS101405.

## Key Findings

- Increased severity of closed head injury in adult male rats is associated with enhanced headache-like behaviors when compared to data obtained in a previous study of a milder injury.
- Repetitive, non-concussive mild closed head injury gives rise to enhanced headache-like behaviors similar to those precipitated by a more severe injury.
- Extended headache-like behaviors following severe, and repetitive mild closed head injury are ameliorated by early and prolonged anti-CGRP mAb treatment, suggesting a mechanism linked to peripheral CGRP signaling.

